# Bundle-specific associations between white matter microstructure and Aβ and tau pathology at their connecting cortical endpoints in older adults at risk of Alzheimer’s disease

**DOI:** 10.1101/2020.08.27.266551

**Authors:** Alexa Pichet Binette, Guillaume Theaud, François Rheault, Maggie Roy, D. Louis Collins, John C.S. Breitner, Judes Poirier, Maxime Descoteaux, Sylvia Villeneuve, for the PREVENT-AD Research Group

## Abstract

Beta-amyloid (Aβ) and tau proteins, the pathological hallmarks of Alzheimer’s disease (AD), are believed to spread through connected regions. Combining diffusion imaging and positron emission tomography, we investigated associations between Aβ, tau and white matter microstructure specifically in bundles connecting brain regions in which AD pathology accumulates. In 126 cognitively normal elderly at risk of AD, we focussed on free-water corrected diffusion measures in the cingulum, posterior cingulum, fornix and uncinate fasciculus. We found higher tissue fractional anisotropy and lower mean and radial diffusivity related to increased Aβ at the cortical endpoints of the cingulum and fornix. We observed similar but stronger associations in the uncinate fasciculus, but with increased Aβ and tau at the endpoints of this bundle. This consistent pattern of associations, with opposite directionality to the usual degeneration pattern in symptomatic individuals, suggests more restricted diffusion in bundles vulnerable to preclinical AD pathology.

## 1. Introduction

The progression of Alzheimer’s disease (AD) neurodegeneration includes a long asymptomatic phase, during which accumulating pathology is accompanied by various brain changes (Jack et al., 2013; Sperling et al., 2011). Beta-amyloid (Aβ) and tau proteins, the pathological hallmarks of the disease (Duyckaerts et al., 2009), start to accumulate decades before signs of cognitive impairment (Bateman et al., 2012; Jansen et al., 2015). Positron emission tomography (PET) can image both proteins *in vivo* (Johnson et al., 2016; Klunk et al., 2004; Schöll et al., 2016), and thus help identifying the earliest brain changes associated with such pathologies. Both Aβ and tau accumulate in a distinct pattern of deposition that follows canonical brain networks/organization. Aβ develops a widespread pattern of deposition that recapitulates a default mode network-like pattern, accumulating early in the frontal and parietal lobes (Mattsson et al., 2019; Villeneuve et al., 2015). Tau accumulates in a more localized pattern, starting in the medial temporal lobe in the preclinical phase of the disease, then spreading later to other parts of the temporal lobe and the rest of the brain in late stages (Braak and Braak, 1991). A prominent view is that pathology accumulates in functionally and/or structurally connected regions (Franzmeier et al., 2019; Seeley et al., 2009; Sepulcre et al., 2017; Vogel et al., 2020). Many studies have highlighted associations between AD pathology and brain functional activity early in the course of the disease (Berron et al., 2020; Jones et al., 2017; Mormino et al., 2011; Sepulcre et al., 2017). However, relations between pathology and white matter measures, as assessed by diffusion magnetic resonance imaging (MRI), remain elusive in preclinical AD. While white matter degeneration is clearly apparent in the late symptomatic stages, how white matter microstructure is affected early on in the disease process is less clear (Sachdev et al., 2013). Whole-brain diffusion MRI tractograms can represent the brain’s white matter architecture, but these are difficult to reconstruct because of extensive crossing of whiter matter fibers and the complexity of tracking algorithms (Rheault et al., 2020). Recent advances in modeling and available algorithms have facilitated robust extraction of white matter bundles with automated methods, thereby allowing their more precise investigation. As well, more specific measures have become available for analysis of white matter (Dyrby et al., 2014). In particular, free-water corrected diffusion tensor measures may offer better estimates of white matter microstructure, yielding tissue-based fractional anisotropy and diffusivities after removing the free-water contribution to each voxel (Pasternak et al., 2009).

In 126 cognitively normal older adults at increased risk of AD, we investigated various diffusion-based measures of white matter microstructure in bundles that connect cortical regions vulnerable to Aβ and tau deposition. As both of these pathologic proteins are thought to accumulate in connected regions, we hypothesized that diffusion measures in white matter bundles would first associate with the amount of pathology specifically in grey matter areas connected by such bundles rather than with more global measures of pathology. We sought to expand upon the few studies linking preclinical AD pathology and white matter microstructure and focussed on *a priori* bundles connecting brain regions targeted early by AD pathology, notably the cingulum bundle (Jacobs et al., 2018). The latter is a large association bundle under the cingulate gyri that connects anterior to posterior cingulate regions and curves further into the parahippocampal gyri of the temporal lobe. This bundle is typically affected in symptomatic AD dementia (Bubb et al., 2018; Jacobs et al., 2018; Kantarci et al., 2017; Roy et al., 2020; Wen et al., 2019) and given its location, could be preferentially affected by Aβ, particularly in its anterior segment. Also of interest is the uncinate fasciculus, reported to be affected at the stage of mild cognitive impairment (Mito et al., 2018; Roy et al., 2020). This bundle connects parts of the limbic system, such as the hippocampus and amygdala in the temporal lobe, with the orbitofrontal cortex (Von Der Heide et al., 2013), brain regions thought to be key regions for tau and Aβ propagation respectively (van der Kant et al., 2020). Lastly, the fornix, which originates in the hippocampus, is another key bundle for investigation that could be affected by tau (Oishi and Lyketsos, 2014; Strain et al., 2018).

## 2. Results

### 2.1 Approach and participantsxs

Using state-of-the-art methods in diffusion MRI modeling, tractography and tractometry, we aimed to better understand the associations between white matter microstructure of key bundles in preclinical AD and deposition of Aβ and tau at their endpoints. We reasoned that the preclinical stage of AD should be the ideal point at which to study these questions, given that this is a period during which AD pathology is spreading but overall brain structure and function remain largely preserved. We therefore studied a subset of 126 asymptomatic individuals at high risk of AD dementia from the PREVENT-AD cohort (Breitner et al., 2016). This cohort enrols cognitively normal older adults at risk of sporadic AD given their parental or multiple-sibling family history of the disease. At time of study, participants were on average 67.3 years of age, predominantly female and highly educated (Table 1). Based on a threshold established previously using global cortical Aβ burden (McSweeney et al., 2020), we estimated that 20% of the participants would be considered Aβ-positive. All underwent diffusion MRI an average of 1.1 ± 0.8 years prior to PET imaging (one completed MRI 5 years prior to PET, but results were unchanged when this participant was removed from analyses).

**Table 1.** Demographics.

|  | **PREVENT-AD participants (n=126)** |
| --- | --- |
| **Age (years)** | $67.3 \pm 4.8$ (68.6-83.2) |
| **Sex F:M (%F)** | 94:32 (75%) |
| **APOE4 carriers (%)** | 50 (40%) |
| **Education (years)** | $15.2 \pm 3.3$ (7.0-24.0) |
| **Global Aβ SUVR** | $1.33 \pm 0.33$ (1.0-2.8) |
| **Meta-ROI temporal tau SUVR** | $1.18 \pm 0.12$ (0.87-2.0) |
| **Mini-Mental State Examination** | $28.8 \pm 1.2$ (24-30) |

Values represent Mean ± SD (Range). Participants with at least one $\epsilon$ 4 allele were considered APOE4 positive. The Mini-mental state evaluation was administered at the same time as PET.
Aβ : beta-amyloid; APOE: apolipoprotein E; SUVR: standardized uptake value ratio

### 2.2. Methodology overview

We extracted free-water corrected diffusion tensor measures and fiber orientation distribution function (fODF)-based measures in bundles of interest. We reconstructed each individual’s whole-brain tractogram using high angular resolution diffusion imaging and fODF, and employed automated tools to isolate the cingulum, the posterior cingulum, the uncinate fasciculus and the fornix (Garyfallidis et al., 2018; Rheault et al., 2018). Tractometry then generated bundle-specific quantification of six white matter properties (Cousineau et al., 2017; Rheault et al., 2017). These were tissue fractional anisotropy (FA_T_), mean diffusivity (MD_T_), axial diffusivity (AD_T_), and radial diffusivity (RD_T_). In each, ‘T’ represents tissue in these free-water corrected diffusion tensor measures. We also report the free-water (FW) index, and fixel-based apparent fiber density (AFD). Free-water is thought to indicate a measure of neuroinflammation (Pasternak et al., 2009). AFD is thought to be more sensitive than FA and an indirect measure of axonal degeneration as it reflects the *apparent* number of axons (Raffelt et al., 2012). We calculated AFD specific to the *fiber population* within a single *voxel*, as per a *fixel-*based approach (Raffelt et al., 2015). To investigate the local relationships with AD pathology, we measured the Aβ and tau levels specifically at the cortical endpoints of each bundle. This approach allowed direct comparisons of the bundles of interest and associated pathology in their connected grey matter regions. An overview of the processing steps is shown in Figure 1. We further evaluated whether associations were independent of atrophy in connected cortical regions, and whether similar associations could be detected with typical diffusion tensor measures, i.e. FA, MD, AD, and RD (not corrected for free-water). Finally, we repeated the main analyses using a global measure of Aβ and a temporal lobe measure of tau instead of testing for associations with AD pathology in grey matter areas connected by the respective white matter bundles. Our premise was that, if AD pathology propagates in connected regions, the associations between pathology and white matter measures should be evident specifically at the endpoints of these bundles.

**Figure 1.**
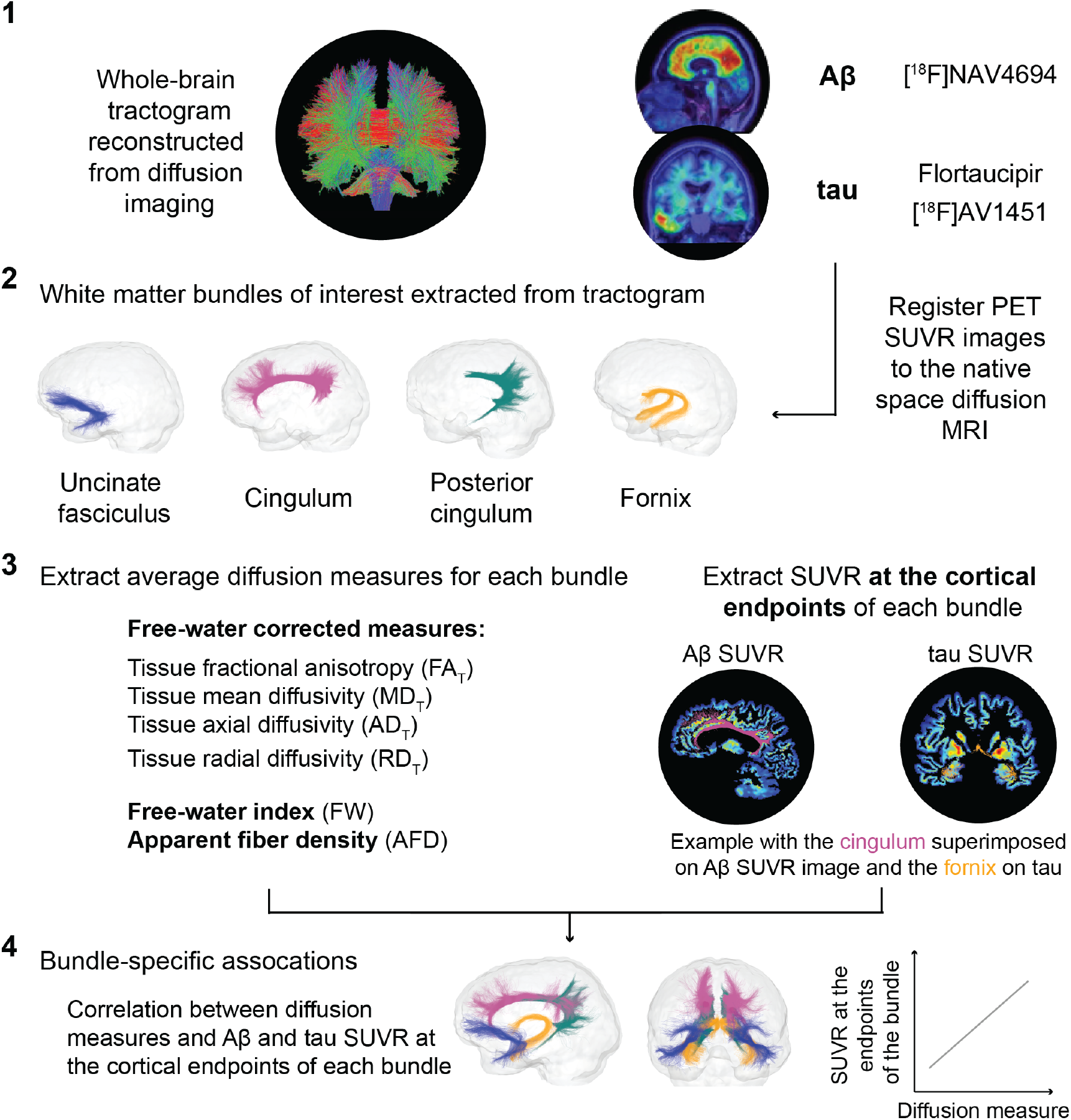
Overview of the processing steps. Whole-brain tractogram reconstructed using the TractoFlow Atlas-Based Segmentation pipeline and automated bundle extraction tools used to extract the four bundles of interest. Free-water corrected tensor measures and total apparent fiber density were calculated for each bundle. PET images were registered to the diffusion space in order to extract the Aβ and tau SUVR directly at the cortical endpoints of each bundle. Aβ: beta-amyloid; PET: positron emission tomography; SUVR: standardized uptake value ratio

### 2.3. Associations in the cingulum and the fornix restricted to Aβ

Examining the amount of pathology at the cortical endpoints along the cingulum, and in the temporal lobe with the fornix, we detected a similar pattern of association with white matter measures and Aβ in both bundles. More specifically, in the right anterior cingulum and right fornix, higher FA_T_, lower MD_T_, and lower RD_T_ were related to higher Aβ endpoint SUVR (Figure 2, Table 2 for the cingulum and Table 3 for the fornix). To evaluate whether such associations with Aβ were independent of tau pathology, we added tau SUVR at the corresponding endpoints as a covariate in these models. We found increased associations with endpoint Aβ in both bundles after adjusting for tau (Table 2 and 3). To evaluate whether association with Aβ was also affected by atrophy in endpoint brain regions, we added grey matter volume of the following: anterior or posterior cingulate cortex in models with the cingulum; and hippocampal volume in models with the fornix. In both instances, associations increased with inclusion of terms for grey matter volume (Table 2 and 3). Note that for the cingulum, however, that this was true only with respect to volume of the anterior cingulate cortex. We found no associations between any white matter measures and tau in either the cingulum or the fornix (Supplementary Figure 1). Examining the same relations in the posterior cingulum, we found no associations between any diffusion measure and either Aβ or tau in this bundle (Supplementary Figure 2).

**Figure 2.**
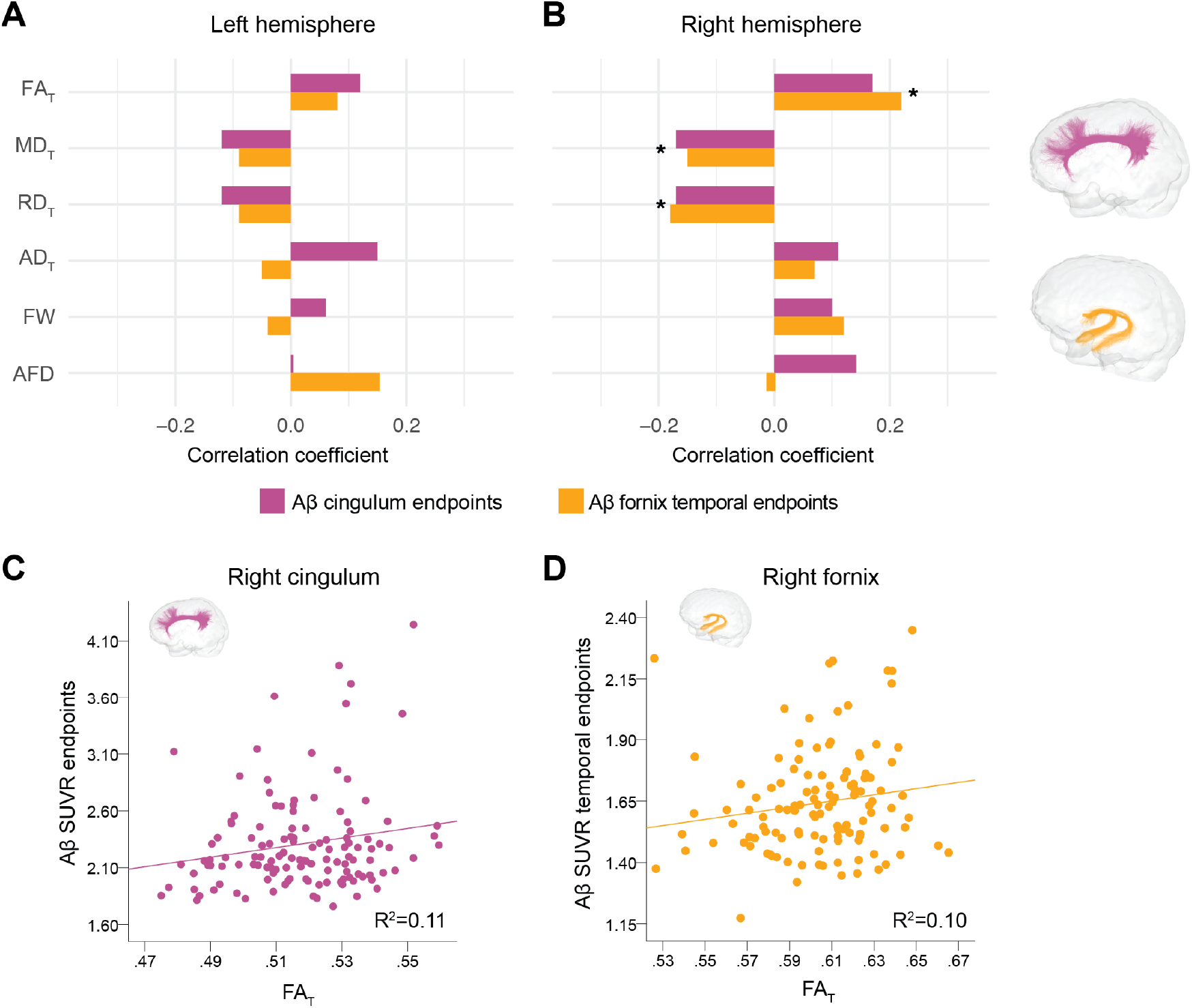
Associations between diffusion measures and Aβ in the cingulum and temporal fornix. R_partial_ from regression models investigating associations between each diffusion measure (average diffusion measure in the bundle; independent variable) and Aβ pathology at the cortical endpoints of the bundle in the left (A) and right (B) hemispheres. Magenta bars correspond to associations in the cingulum and orange bars, in the fornix. Models included age, sex, bundle volume (divided by total intracranial volume) as covariates. * represents consistent associations at p<0.05 for each bundle when further adjusting for either tau pathology or grey matter volume. Associations between FA_T_ and Aβ in the right cingulum (C) and right fornix (D) Aβ: beta-amyloid; FA_T_: tissue fractional anisotropy; MD_T_: tissue mean diffusivity; AD_T_: tissue axial diffusivity; RD_T_: tissue radial diffusivity; FW: free-water index; AFD: fixel-based apparent fiber density

**Table 2.** Associations between diffusion measures and pathology in the cingulum.

| n=126 | Left hemisphere |  |  |  |  |  | Right hemisphere |  |  |  |  |  |
| --- | --- | --- | --- | --- | --- | --- | --- | --- | --- | --- | --- | --- |
| Associations with Aβ |  |  |  |  |  |  |  |  |  |  |  |  |
|  | FA<sub>T</sub> | MD<sub>T</sub> | AD<sub>T</sub> | RD<sub>T</sub> | FW | AFD | FA<sub>T</sub> | MD<sub>T</sub> | AD<sub>T</sub> | RD<sub>T</sub> | FW | AFD |
| Cingulum endpoints | 0.12<br>(0.20) | -0.12<br>(0.20) | 0.15<br>(0.11) | -0.12<br>(0.20) | 0.06<br>(0.55) | 0.00<br>(0.99) | 0.17<br>(0.06)<br>*,<sup>1</sup> | -0.17<br>(0.06)<br>*,<sup>1</sup> | 0.11<br>(0.25) | -0.17<br>(0.06)<br>*,<sup>1</sup> | 0.10<br>(0.30) | 0.13<br>(0.17) |
| Associations with tau |  |  |  |  |  |  |  |  |  |  |  |  |
| Cingulum endpoints | 0.05<br>(0.58) | -0.05<br>(0.58) | 0.08<br>(0.36) | -0.05<br>(0.57) | 0.06<br>(0.50) | 0.05<br>(0.60) | -0.05<br>(0.61) | 0.05<br>(0.61) | 0.00<br>(0.97) | 0.05<br>(0.61) | 0.00<br>(0.98) | 0.07<br>(0.47) |

R<sub>partial</sub> and (p-values) from regression models investigating associations between each diffusion measure (average diffusion measure in the bundle; independent variable) and pathology at the cortical endpoints along the cingulum (dependent variable). Models included age, sex, bundle volume (divided by total intracranial volume) as covariates.
\* becomes significant (p=0.04) when adding tau endpoints SUVR as a covariate
<sup>1</sup> becomes significant (p=0.02) when adding anterior cingulate volume (divided by total intracranial volume) as a covariate, but remains unchanged when adding posterior cingulate volume as a covariate
<sup>2</sup> becomes significant (p=0.04) when adding anterior cingulate volume as a covariate
Aβ : beta-amyloid; FA<sub>T</sub>: tissue fractional anisotropy; MD<sub>T</sub>: tissue mean diffusivity; AD<sub>T</sub>: tissue axial diffusivity; RD<sub>T</sub>: tissue radial diffusivity; FW: free-water index; AFD: fixel-based apparent fiber density

**Table 3.** Associations between diffusion measures and pathology in the fornix.

| Left hemisphere (n=118) |  |  |  |  |  |  | Right hemisphere (n=120) |  |  |  |  |  |
| --- | --- | --- | --- | --- | --- | --- | --- | --- | --- | --- | --- | --- |
| Associations with Aβ |  |  |  |  |  |  |  |  |  |  |  |  |
|  | FA<sub>T</sub> | MD<sub>T</sub> | AD<sub>T</sub> | RD<sub>T</sub> | FW | AFD | FA<sub>T</sub> | MD<sub>T</sub> | AD<sub>T</sub> | RD<sub>T</sub> | FW | AFD |
| Temporal endpoints | 0.08<br>(0.38) | -0.09<br>(0.35) | -0.05<br>(0.62) | -0.09<br>(0.34) | -0.04<br>(0.67) | 0.15<br>(0.13) | 0.22<br>(0.02)<br>*,<sup>1</sup> | -0.15<br>(0.11)<br>*,<sup>1</sup> | 0.07<br>(0.48) | -0.18<br>(0.06)<br>*,<sup>1</sup> | 0.12<br>(0.19) | -0.01<br>(0.92) |
| Associations with tau |  |  |  |  |  |  |  |  |  |  |  |  |
| Temporal endpoints | -0.15<br>(0.12) | 0.16<br>(0.09) | 0.08<br>(0.38) | 0.16<br>(0.09) | -0.10<br>(0.29) | 0.02<br>(0.88) | -0.10<br>(0.27) | 0.05<br>(0.61) | -0.08<br>(0.40) | 0.07<br>(0.46) | -0.01<br>(0.88) | -0.06<br>(0.5) |

R<sub>partial</sub> and (p-values) from regression models investigating associations between each diffusion measure (average diffusion measure in the bundle; independent variable) and pathology at the temporal endpoints of the fornix (dependent variable). Models included age, sex, bundle volume (divided by total intracranial volume) as covariates.
\* more variance was explained when adding tau endpoints SUVR as a covariate (R=0.27, p=0.003 for FA<sub>T</sub>; R=-0.18, p=0.05 for MD<sub>T</sub>; R=-0.22, p=0.02 for RD<sub>T</sub>)
<sup>1</sup> more variance was explained when adding hippocampal volume (divided by total intracranial volume) as a covariate (R=0.23, p=0.01 for FA<sub>T</sub>; R=-0.18, p=0.05 for MD<sub>T</sub>; R=-0.21, p=0.03 for RD<sub>T</sub>)
Aβ : beta-amyloid; FA<sub>T</sub>: tissue fractional anisotropy; MD<sub>T</sub>: tissue mean diffusivity; AD<sub>T</sub>: tissue axial diffusivity; RD<sub>T</sub>: tissue radial diffusivity; FW: free-water index; AFD: fixel-based apparent fiber density

### 2.4. Associations in the uncinate fasciculus with Aβ and tau

Overall, a consistent pattern of associations between white matter microstructure and pathology was apparent across the uncinate fasciculus, anterior cingulum and fornix. The strongest relations between AD pathology and diffusion measures were observed in the uncinate fasciculus. Associations with both Aβ and tau were detected bilaterally at both its endpoints, i.e. in the frontal and temporal lobes (Figure 3, Table 4). More specifically, as in the cingulum and the fornix, higher FA_T_, lower MD_T_, lower RD_T_, and higher AFD related to higher Aβ (Figure 3A to C) and tau (Figure 3D to F) SUVR at bundle endpoints. Higher AD_T_ also related to higher tau SUVR, and higher FW index related to higher tau endpoint SUVR for the right uncinate fasciculus only (Figure 3D). All associations with frontal tau endpoint SUVR survive correction for multiple comparisons (Table 4). To evaluate whether associations detected with Aβ or tau pathology were independent, we added tau SUVR in corresponding endpoints as a covariate in models with Aβ as the dependent variable, and vice versa. The original significant associations with Aβ appeared slightly less impressive (p=0.08) with insertion of the tau covariates (Supplementary Table 1). However, initial associations with tau remained after inserting covariates for Aβ at bundle endpoints (Supplementary Table 1). Lastly, we added grey matter volume from the medial orbitofrontal cortex and from the parahippocampal gyri as a covariate in models assessing frontal and temporal endpoint SUVRs, respectively (Table 4). Apparently, atrophy has little influence on associations between white matter microstructure and pathology in this bundle as all associations remained.

**Figure 3.**
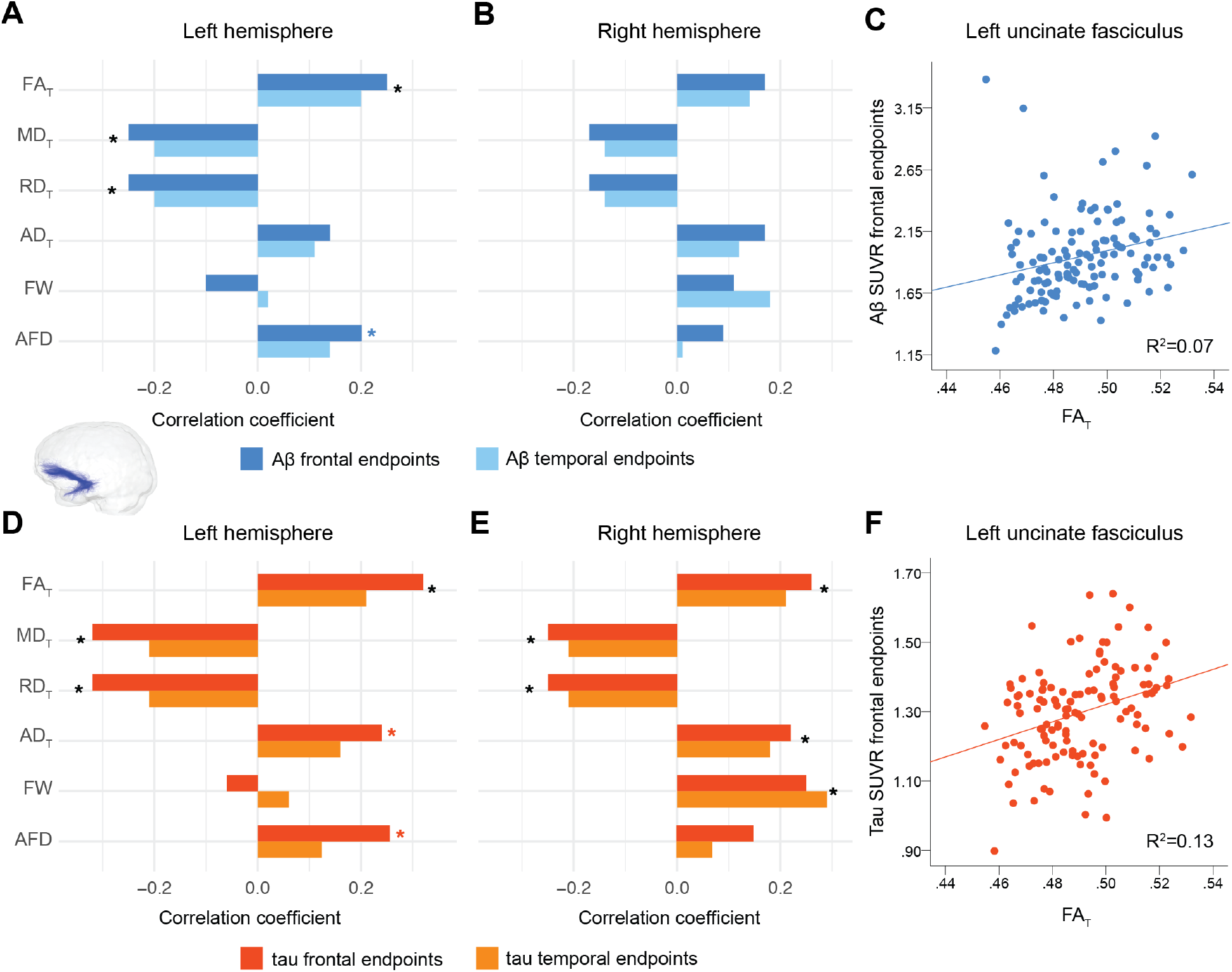
Associations between diffusion measures and pathology in the uncinate fasciculus. R_partial_ from regression models investigating associations between each diffusion measure (average diffusion measure in the bundle; independent variable) and Aβ (A-B) or tau (D-E) pathology at the endpoints in the two ends of the uncinate fasciculus in the left and right hemispheres. Models included age, sex, bundle volume (divided by total intracranial volume) as covariates. Representative associations between FA_T_ with Aβ and tau are shown in C and F. * represents p<0.05 and is black when associations at both endpoints were significant, and red when association at the frontal endpoints only was significant. Aβ: beta-amyloid; FA_T_: tissue fractional anisotropy; MD_T_: tissue mean diffusivity; AD_T_: tissue axial diffusivity; RD_T_: tissue radial diffusivity; FW: free-water index; AFD: fixel-based apparent fiber density

**Table 4.** Associations between diffusion measures and pathology in the uncinate.

| n=126 | Left hemisphere |  |  |  |  |  | Right hemisphere |  |  |  |  |  |
| --- | --- | --- | --- | --- | --- | --- | --- | --- | --- | --- | --- | --- |
| Associations with Aβ |  |  |  |  |  |  |  |  |  |  |  |  |
|  | FA<sub>T</sub> | MD<sub>T</sub> | AD<sub>T</sub> | RD<sub>T</sub> | FW | AFD | FA<sub>T</sub> | MD<sub>T</sub> | AD<sub>T</sub> | RD<sub>T</sub> | FW | AFD |
| Frontal endpoints | 0.25<br>(**0.005**)<sup>*</sup> | -0.25<br>(**0.005**)<sup>*</sup> | 0.14<br>(0.12) | -0.25<br>(**0.005**)<sup>*</sup> | -0.10<br>(0.28) | 0.20<br>(0.026)<sup>*</sup> | 0.17<br>(0.06)<sup>+</sup> | -0.17<br>(0.06)<sup>+</sup> | 0.17<br>(0.07)<sup>+</sup> | -0.17<br>(0.06)<sup>+</sup> | 0.11<br>(0.21) | 0.08<br>(0.37) |
| Temporal endpoints | 0.20<br>(0.03)<sup>*</sup> | -0.20<br>(0.03)<sup>*</sup> | 0.12<br>(0.20) | -0.20<br>(0.03)<sup>*</sup> | 0.02<br>(0.84) | 0.13<br>(0.15) | 0.14<br>(0.14) | -0.14<br>(0.14) | 0.12<br>(0.20) | -0.14<br>(0.14) | 0.18<br>(0.05) | 0.01<br>(0.88) |
| Associations with tau |  |  |  |  |  |  |  |  |  |  |  |  |
| Frontal endpoints | 0.32<br>(**<0.001**)<sup>*</sup> | -0.32<br>(**<0.001**)<sup>*</sup> | 0.24<br>(**0.01**)<sup>*</sup> | -0.32<br>(**<0.0001**)<sup>*</sup> | -0.06<br>(0.51) | 0.24<br>(**0.008**)<sup>*</sup> | 0.26<br>(**0.005**)<sup>*</sup> | -0.25<br>(**0.005**)<sup>*</sup> | 0.22<br>(**0.01**)<sup>*</sup> | -0.25<br>(**0.005**)<sup>*</sup> | 0.25<br>(**0.006**)<sup>*</sup> | 0.15<br>(0.10) |
| Temporal endpoints | 0.21<br>(0.02)<sup>*</sup> | -0.21<br>(0.02)<sup>*</sup> | 0.16<br>(0.08) | -0.21<br>(0.02)<sup>*</sup> | 0.06<br>(0.54) | 0.11<br>(0.24) | 0.21<br>(0.02)<sup>*</sup> | -0.21<br>(0.02)<sup>*</sup> | 0.18<br>(0.05)<sup>*</sup> | -0.21<br>(0.02)<sup>*</sup> | 0.29<br>(0.001)<sup>*</sup> | 0.06<br>(0.54) |

$R_{\text{partial}}$ and (p-values) from regression models investigating associations between each diffusion measure (average diffusion measure in the bundle; independent variable) and pathology at the cortical endpoints of the two ends of the uncinate fasciculus (dependent variable). Models included age, sex, bundle volume (divided by total intracranial volume) as covariates. Bolded associations survived FDR correction at $q=0.05$ .
\* association remains unchanged when grey matter volumes (divided by total intracranial volume) at the endpoint of the bundles were added as covariates in the respective models.
+ association becomes significant ( $p=0.05$ ) when adding medial orbitofrontal cortex volume as a covariate
Aβ : beta-amyloid; FA<sub>T</sub>: tissue fractional anisotropy; MD<sub>T</sub>: tissue mean diffusivity; AD<sub>T</sub>: tissue axial diffusivity; RD<sub>T</sub>: tissue radial diffusivity; FW: free-water index; AFD: fixel-based apparent fiber density

### 2.5 Importance of advanced free-water measures to these results

To evaluate the sensitivity of free-water corrected measures over the typical tensor measures, we tested whether similar associations with pathology exist with FA, MD, AD and RD (i.e. not corrected for free-water). We failed to observe most of the above-described associations when using these classical diffusion tensor measures. Specifically, no associations were found between Aβ or tau and any of the diffusion tensor measures in the cingulum or posterior cingulum. In the fornix, there was one isolated relationship between FA and the left temporal Aβ endpoints (R_partial_=0.24; p=0.01). In the uncinate fasciculus, only a few associations were detected, the main ones being higher FA and AD related to higher tau pathology (Supplementary Table 2). The directionality of the associations was the same as for the free-water corrected measures.

### 2.6 No associations with global measures of Aβ or tau pathology

Lastly, we found almost no associations between either Aβ or tau global scores and the diffusion measures in any of the bundles using either free-water corrected or classical tensor measures. The sole exception was one isolated association of higher FW index in the right uncinate fasciculus and higher tau SUVR, with R_partial_=0.23 (p=0.01).

## 3 Discussion

The notion that AD pathology spreads through connected regions in the brain has foundations in rodent models (Ahmed et al., 2014; Palop and Mucke, 2010), although it is gaining credence in human neuroimaging studies. The spatial overlap between key regions of Aβ accumulation and connectivity hubs is striking (Buckner et al., 2005; Sepulcre et al., 2016). Tau has been found to propagate in connected regions independently of distance between them (Franzmeier et al., 2020; Vogel et al., 2020). Bundle-specific white matter neurodegeneration has also been associated with increased tau accumulation (Jacobs et al., 2018). Combining Aβ- and tau-PET with recent advanced diffusion imaging analyses, we investigated AD pathology and white matter microstructure in *a priori* selected bundles that connect the brain regions where pathology accumulates. Our aim here was not to test the spreading hypothesis *per se* but, assuming that this hypothesis is correct, to focus on local effects of white matter bundle microstructure and pathology to increase probabilities of capturing early alterations. To do so, we investigated microstructure – pathological associations in a cohort of asymptomatic older adults younger than most typical aging or AD cohorts and enriched for preclinical AD due to their family history of the disease. We characterized the association focussing at a macro-level on key bundles of interest, and at a micro-level on diffusion measures in these bundles and their associations with Aβ and tau levels specifically at their cortical endpoints. We found a consistent pattern of more restricted diffusion across the cingulum, the fornix, and the uncinate fasciculus, with higher FA_T_, lower MD_T_, and lower RD_T_ being related to greater endpoint pathology. For the cingulum and fornix such associations were restricted to Aβ, but they were found both with Aβ and tau in the uncinate fasciculus.

There are two important take-homes related to the main results. First, the pattern of association between pathology and microstructure was only detected using local Aβ and tau at the bundle endpoints and not when using global AD burden scores. This finding suggests topographical relationships between pathology and white matter microstructural alterations in the early stage of AD. This “bundle-specific” approach through tractography and tractometry complements the typical approach of voxel-wise analyses (Harrison et al., 2020; Zhang et al., 2019) and yielded specific associations between white matter and pathology. Similarly, using more precise tissue measures with free-water corrected as opposed to classical diffusion tensor measures was critical to our findings, further highlighting the relevance of novel methods. Second, the directionality of the observed pattern of association opposes the classical pattern of degeneration. The classical degeneration pattern accompanying disease progression is characterized by lower anisotropy and higher diffusivity, representing loss of coherence in the white matter microstructure with AD progression (Caso et al., 2016; Sexton et al., 2011). This pattern of white matter degeneration develops invariably along the AD spectrum (Amlien and Fjell, 2014; Pereira et al., 2019), with associations often becoming detectable only in the mild cognitive impairment and dementia stages (Mito et al., 2018; Song et al., 2018; Wang et al., 2019; Wen et al., 2019), and very few in Aβ-positive cognitively normal participants (Rieckmann et al., 2016; Vipin et al., 2019). However, our consistent pattern of more restricted diffusion (higher FA and lower MD) being associated with more pathology in three bundles suggests that microstructural alterations captured with diffusion MRI might differ in the preclinical vs. the symptomatic phase of AD, during which severe and irreversible atrophy has occurred. Supporting this biphasic relationship, in the Alzheimer Disease Neuroimaging Initiative dataset, FA increased from the Aβ-negative participants to those with intermediate Aβ levels and decreased in those with high Aβ (Dong et al., 2020). In another cohort of older adults with intact cognition and a family history of AD, those with high levels of Aβ also had higher FA and lower MD compared to the Aβ-negative participants, notably in the cingulum and the fornix (Racine et al., 2014). This possible biphasic relationship (Fortea et al., 2010; Montal et al., 2018) has important implication as it might obscure some association in the early disease stages.

Although more restricted diffusion with the presence of pathology was unanticipated, different biological mechanisms might underlie this phenomenon. For example, a loss of crossing fibers occurring as pathology accumulates could account for increased FA_T_ in the early disease stage (Mito et al., 2018). Further, higher anisotropy and lower diffusivity might be due to hypertrophy, glial activation, neuronal or glial swelling in the asymptomatic phase (Fortea et al., 2010; Montal et al., 2018). Extracellular Aβ plaques typically accumulate in the grey matter but have often been associated with gliosis (Spires-Jones and Hyman, 2014), which could restrict diffusion in the microstructural environment, as reported here with higher FA_T_ and lower MD_T_ and RD_T_. Tau, on the other hand, is a microtubule-associated protein that stabilizes the axons. With disease progression, tau becomes hyperphosphorylated and detaches from the microtubules (Higuchi et al., 2002; Iqbal et al., 2009). We can hypothesize that the increasing presence of hyperphosphorylated tau along axons early on in AD pathological processes could restrict the diffusion, even more so when looking at tissue measures specifically.

We were surprised to find no associations between AD pathology and diffusion measures in the posterior cingulum, as this is a key bundle in AD (Agosta et al., 2011; Caso et al., 2016; Zhuang et al., 2012). This bundle is certainly altered in the symptomatic stage, but it is possible that the microstructure of the posterior part of the cingulum is not affected early by pathology. Alternatively, this region might have already entered a shift toward the classical degeneration pattern (lower FA_T_ and AFD, higher MD_T_ and RD_T_) that would be related to higher AD pathology in some individuals. This opposite direction of associations that switches with disease severity would make linear findings impossible to detect in individuals that are slowly progressing from cognitively normal to cognitively impaired. On the other hand, the strongest and more numerous associations were detected in the uncinate fasciculus. This bundle has an interesting anatomy, connecting regions at the intersection of both Aβ (frontal lobe) and tau (temporal lobe) deposition patterns. We speculate that the particular localization of the uncinate fasciculus with regards to Aβ and tau deposition might confer early vulnerability to pathological insults. Its possible vulnerability might also be increased due to retrogenesis. This concept postulates that late-myelinated fibers, from temporal and neocortical regions, are affected first in the disease course, whereas thicker fibers myelinated earlier in development are more resistant to neurodegeneration/disease (Alves et al., 2015; Bartzokis, 2004, 2011). For instance, the orbitofrontal cortex is not only a region where Aβ pathology accumulates early but is also a highly plastic late-developing region, typically affected in aging (Fjell et al., 2014; Pichet Binette et al., 2020).

The direct investigation of pathology at the cortical endpoints of white matter fiber bundles and microstructure in such bundles was possible due to recent advances in diffusion imaging modeling, tractography, bundle extraction and tractometry quantification. However, there are several limitations to these techniques and to our study. First, there are no common standards (yet) to extract pre-defined bundles from tractograms, and bundles with high curvature such as the uncinate fasciculus and the fornix are challenging to extract. To mitigate this challenge, we mostly relied on algorithms that use priors to help generate fuller bundles. Still, to extract all bundles of interest, we needed to use multiple automated algorithms and perform rigorous visual inspection to make sure all algorithms yielded comparable bundles. The diffusion sequence relied on only one b-value, and future acquisitions with multiple b-values could further improve capturing fine-grained changes (Pines et al., 2020). Given the partial volume effect of PET, pathology at the cortical endpoints might be slightly affected with white matter uptake. To diminish this potential confound, we took advantage of the high 2-mm resolution of the scanner (HRRT) and did not smooth the SUVR images. This cohort is followed yearly on cognition and imaging, so future longitudinal studies will help clarify the potential biphasic relationship, as a proportion of participants will progress to mild cognitive impairment.

Overall, we used state-of-the-art analytical techniques to study associations between white matter microstructure and pathology in key bundles affected in AD in the PREVENT-AD cohort of cognitively normal older adults whose strong family history of AD suggests a two- to three-fold increased risk of subsequent dementia (Cupples et al., 2004; Devi et al., 2000). We suggest that our reliance on this cohort was important because AD pathology starts depositing in the asymptomatic phase of the disease, but extensive cortical pathology and atrophy are apparent by the time an individual develops cognitive impairment. PREVENT-AD and similar samples represent the sorts of groups that may be useful for clinical trials of preventive interventions (Meyer et al., 2019). As more studies highlight that white matter changes might precede changes in grey matter (Caso et al., 2016; Sachdev et al., 2013), studying the associations between pathology and microstructure in the early stages of AD will help understand better of the complex pathogenesis of the disease.

## 4 Materials and Methods

### 4.5 Participants

We studied cognitively unimpaired participants at risk of sporadic AD dementia from the PRe-symptomatic EValuation of Experimental or Novel Treatments for AD (PREVENT-AD) study. PREVENT-AD is a longitudinal study that started in 2012 (Breitner et al., 2016) and enrolled 386 participants. Inclusion criteria were as follows: (1) having intact cognition, (2) having a parent or two siblings diagnosed with AD-like dementia, and therefore being at increased risk of sporadic AD, (3) being above 60 years of age, or between 55 and 59 if fewer than 15 years from their affected family member’s age at symptom onset, (4) being free of major neurological and psychiatric diseases. Intact cognition was based on the Montreal Cognitive Assessment, a Clinical Dementia Rating of 0, and a standardized neuropsychological evaluation using the Repeatable Battery for the Assessment of Neuropsychological Status (Randolph et al., 1998). The cognitive status of individuals with questionable neuropsychological status was reviewed in consensus meetings of neuropsychologists (including SV) and/or psychiatrists. Annual visits include neuropsychological testing and a MRI session. Since 2017, Aβ and tau PET scans were integrated to the study protocol for interested participants. The present study includes participants who had structural and diffusion-weighted MRI and who underwent PET, for a total of 126 participants. All participants included in the current study were cognitively normal at the time they underwent diffusion-weighted MRI.

### 4.6 Image Acquisition

#### 4.6.1 Magnetic resonance imaging

T1-weighted structural and diffusion-weighted MRI were acquired on a Magnetom Tim Trio 3 Tesla (Siemens) scanner at the Douglas Mental Health University Institute prior to PET imaging. Structural scans were acquired yearly, and thus we selected the closest scan prior to PET (average time between PET and structural MRI: 8 ± 4 months). Diffusion-weighted MRI was not acquired every year, and again the diffusion scan closest to PET was chosen for analysis (average time between PET and diffusion-weighted MRI: 1.1 ± 0.8 years). Structural scans were acquired using a MPRAGE sequence with the following parameters: TR=2300 ms; TE =2.98ms; FA=9°; FoV=256 mm; slice thickness=1mm; 160-170 slices. Diffusion-weighted scans were acquired with the following parameters: TR=9300 ms, TE: 92 ms, FoV=130 mm, slice thickness=2 mm. One b0 image was acquired and 64 diffusion-weighted volumes were acquired with a b-value of 1000 s/mm^2^.

### 4.6.2 Positron emission tomography

PET was performed using [^18^F]NAV4694 to assess Aβ burden and flortaucipir ([^18^F]AV1451) to assess tau deposition. PET scanning took place at the McConnell Brain Imaging Centre at the Montreal Neurological Institute using a brain-dedicated PET Siemens/CT high-resolution research tomograph (HRRT) on two consecutive days. Aβ scans were acquired 40 to 70 minutes post-injection (≈6 mCi) and tau scans 80 to 100 minutes post-injection (≈10 mCi). All scans were completed between March 2017 and April 2019.

### 4.7 Positron emission tomography processing

PET scans were processed using a standard pipeline (see https://github.com/villeneuvelab/vlpp for more details). Briefly, Aβ- and *tau*-PET images were realigned, averaged and registered to the T1-weighted scan of each participant, which had been segmented with the Desikan-Killiany atlas using FreeSurfer version 5.3 (Desikan et al., 2006). The same structural scan was used in the diffusion and the PET pipelines. PET images were then masked to remove the scalp and cerebrospinal fluid, to reduce contamination by non-grey and non-white matter voxels. Standardized uptake value ratios (SUVR) images were obtained using the whole cerebellum as reference region for Aβ-PET (Jagust et al., 2015) and the inferior cerebellar grey matter for tau-PET (Baker et al., 2017). A global Aβ burden was calculated from the average bilateral SUVR of medial and lateral frontal, parietal and temporal regions. A global score of temporal tau was calculated by taking the average bilateral SUVR from the entorhinal, amygdala, fusiform, inferior and middle temporal gyri (Ossenkoppele et al., 2018). However, the main interest was to extract Aβ and tau SUVR at the cortical endpoints of the fibers forming each anatomical bundle of interest, and thus we registered the PET SUVR images to the diffusion image using ANTS (Avants et al., 2011). We also took advantage of the high 2-mm resolution of the PET-HRRT scanner and did not smooth the SUVR images. By doing so, we aimed to diminish mixing grey and white matter signal in each voxel, so that the SUVR values at the cortical endpoints of the fibers are more precise.

### 4.8 Diffusion MRI processing

An overview of the processing steps is displayed in Figure 1.

#### 4.8.1 Preprocessing steps

The diffusion-weighted images were processed using the TractoFlow Atlas-Based Segmentation (TractoFlow-ABS) pipeline. TractoFlow-ABS is an extension of the recent TractoFlow pipeline (Theaud et al., 2020a; Theaud et al., 2020b) publicly available for academic research purposes (https://github.com/scilus/tractoflow) that uses Nextflow (Di Tommaso et al., 2017) and Singularity (Kurtzer et al., 2017) to ensure efficient and reproducible diffusion processing. All major processing steps are performed through this pipeline, from preprocessing of the structural and diffusion images to tractography. The pipeline computes typical diffusion tensor imaging maps, fiber orientation distribution function (fODF) and a whole-brain tractogram. The pipeline calls different functions from various neuroimaging software, namely FSL (Jenkinson et al., 2012), MRtrix3 (Tournier et al., 2019), ANTs (Avants et al., 2011), and DIPY (Garyfallidis et al., 2014). For a detailed description of the different steps see (Theaud et al., 2020a).

#### 4.8.2 Diffusion measures

After the preprocessing steps, different diffusion measures can be generated as part of TractoFlow or TractoFlow-ABS. The following diffusion tensor imaging (DTI) metrics were computed using DIPY: fractional anisotropy (FA), mean diffusivity (MD), radial diffusivity (RD) and axial diffusivity (AD). Along with typical DTI modeling, fiber orientation distribution functions (fODFs) were also computed using constrained spherical deconvolution (Descoteaux et al., 2007; Tournier et al., 2007) and the fiber response function from the group average. The fODF metric used in the current study was apparent fiber density along each bundle (AFD), which can be seen as an indirect measure of axonal density (Raffelt et al., 2012). AFD was calculated along each bundle in a fixel-based manner, reducing contamination from crossing fibers (Raffelt et al., 2015).

Along with the typical tensor metrics, we also generated free-water corrected DTI metrics, which were the main diffusion metrics of interest in this study. Free-water correction has been proposed has a way to remove the contamination of water from the tissue properties by modeling the isotropic diffusion of the free water component (Pasternak et al., 2009). Free-water modeling was performed using the accelerated microstructure imaging via convex optimization (Daducci et al., 2015) to calculate free-water index (FW) and free-water corrected metrics, namely FA_T_, MD_T_, AD_T_ and RD_T_. Removing the contribution of free water is thought to better represent the tissue microstructure (hence the subscript _T_ for tissue) and might be more sensitive than the non-corrected metrics (Albi et al., 2017; Chad et al., 2018; Pasternak et al., 2012).

#### 4.8.3 Tractography

The last step of the pipeline is tractography. This is where Tractoflow and Tractoflow-ABS differ. The former uses a more sophisticated algorithm, particle filtering tractography, that takes into account anatomical information to reduce tractography biases (Girard et al., 2014). Such an algorithm requires probabilistic maps of grey matter (GM), white matter (WM) and cerebrospinal fluid to add additional constraints for tracking. However, with aging, probabilistic maps in “bottleneck” areas of WM fibers, for example where the uncinate fasciculus bends, show poorer distinction between GM and WM voxels. Furthermore, increasing white matter hyperintensities and general atrophy with aging also complicate the use of more advanced algorithms. As a result, the performance particle filtering tractography was affected and failed to generate bundles suitable for analysis. Instead, as implemented in TractoFlow-ABS, we opted for local tracking with a probabilistic algorithm to reconstruct whole-brain tractograms. The inputs for tracking were the fODF image for directions and a WM mask for seeding. The mask was computed by joining the WM and the subcortical masks from the structural image that had been segmented with the Desikan-Killiany atlas in FreeSurfer version 5.3 (Desikan et al., 2006). For tracking, seeding was initiated in voxels from the WM mask with 10 seeds per voxel. The tractograms had between 2 and 3 million streamlines.

### 4.9 White matter bundles extraction

From the tractogram, we extracted different bundles of interest. We focussed on bundles connecting the main brain region where Aβ and tau accumulate in the early phase of AD, namely the uncinate fasciculus, the cingulum, the posterior cingulum, and the fornix. To extract the uncinate fasciculus and the cingulum, we used RecoBundles X (Rheault, 2020), an automated algorithm to segment the tractograms into different bundles. This algorithm is an improved and more stable version of RecoBundles (Garyfallidis et al., 2018). Briefly, the method is based on shape priors to detect similarity in streamlines. Taking the whole-brain tractogram and templates from the bundles of interest as inputs, RecoBundles X extracts bundles based on the shape of the streamlines from the templates. The difference between RecoBundles and RecoBundles X resides in that the latter can take multiple templates as inputs and multiple parameters, which refines which streamlines are included or excluded from the final bundle. RecoBundles X is typically run 80 times and the output is the conjunction of the multiple runs, yielding more robust bundles. RecoBundles X does not include templates for the posterior cingulum or the fornix, and thus we used different methods to extract them. We used TractQuerier (Wassermann et al., 2016) to extract the posterior cingulum. This method works with customizable queries to extract bundles based on anatomical definitions. Using inclusion and exclusion regions of interest based on the FreeSurfer parcellation, we implemented a query specifically for the posterior cingulum. The query was also used in another recent study (Roy et al., 2020) and can be found in Supplementary material. The last bundle of interest was the fornix. The fornix is a difficult bundle to extract, given its high curvature, its proximity to cerebrospinal fluid increasing susceptibility to partial volume effects, and its location in regions prone to atrophy in aging and AD,. Therefore, we used a combination of different steps to generate this bundle. First, we ran Bundle-Specific Tractography (Rheault et al., 2019). This algorithm helps to increase the number of plausible streamlines, yielding a better spatial coverage and a more accurate representation of the full shape of the fornix. Bundle-Specific Tractography takes as input a template representing the fornix derived from 23 participants used in another study (Roy et al., 2020). This template, in the form of streamlines, generates spatial and orientational priors to enhance the fODF map and facilitate reconstruction of the fornix. Using Bundle-Specific Tractography to extract the fornix specifically is further detailed elsewhere (Rheault et al., 2018). Applied to cognitively normal older adults such as the PREVENT-AD participants, the algorithm yielded fornices with a very high number of streamlines and we implemented further steps to filter out the fornix tractograms: we excluded any streamlines going through the thalamus and through an eroded mask of CSF, and only retained streamlines going through the hippocampus. Finally, we removed the streamlines forming loops.

After extracting all bundles, each one was inspected visually in MI-Brain (https://www.imeka.ca/fr/mi-brain/) to make sure the shape, location and size were adequate. Only for the fornix did some participants have bundles that were too small. We excluded from analyses fornices containing less than 50 streamlines.

### 4.10 Bundle-specific quantification with tractometry

The last step required to put together the different white matter measures and bundles of interest was tractometry (Cousineau et al., 2017). Tractometry is a way to extract the measures of interest specifically in each bundle. It takes as input the maps of all microstructure measures and the bundles in which we want to extract them. In our case, we extracted the average bundle tissue measures (FA_T_, MD_T_, RD_T_, AD_T_ and FW index) and AFD as a fODF metric for each bundle (uncinate fasciculus, cingulum, posterior cingulum, fornix). For complementary analyses we also extracted typical tensor measures (average FA, MD, RD and AD) in each bundle. For the Aβ and tau measurements, we extracted the average SUVR from the cortical endpoints of each bundle. By doing so, we have an average Aβ/tau SUVR of all voxels specific to each bundle and participant. In the cingulum, the endpoints lie along the cingulate cortex. In the posterior cingulum, we extracted SUVR at both ends of the bundle, i.e. in the posterior cingulate and the medial temporal lobe. In the fornix, we extracted SUVR at the endpoints in the temporal lobe only. We did not consider the endpoints around the mammillary bodies, as they are regions with off-target binding in PET. In the uncinate fasciculus, we extracted SUVR at both ends of the bundle, i.e. in the frontal and temporal lobes. The overall approach, done entirely in native space, has the advantage of generating bundles specific to each individual and of capturing the amount of pathology specifically in the grey matter connected by such bundles.

### 4.11 Statistical Analysis

Linear regression models were performed to evaluate the relationships between Aβ or tau and the different microstructure measures in each bundle. In primary analyses, the diffusion measures investigated as independent variables were FA_T_, MD_T_, RD_T_, AD_T_, FW index and AFD. Regression models were performed separately for Aβ and tau in the left and right bundles separately. Age, sex, and bundle volume (divided by total intracranial volume) were included as covariates in each regression model. We focused on the Aβ and tau SUVR specifically at the endpoints of each bundle. In the bundles where associations were found between pathology and microstructure, we further adjusted for grey matter volume (divided by total intracranial volume) of cortical regions connected by the bundle to evaluate whether associations were also influenced by atrophy. For the uncinate fasciculus, GM regions of interest were medial orbitofrontal cortex and the parahippocampal gyrus; for the cingulum, regions were the anterior and posterior cingulate; for the posterior cingulum, regions were precuneus and parahippocampal gyrus; for the fornix, we used hippocampal volume. Lastly, when we found associations between different microstructure measures and one pathology (Aβ or tau), we further corrected for the SUVR of the other protein at the same endpoints, to assess whether associations were specific to one protein. We also performed similar analyses with the typical tensor measures (FA, MD, AD and RD) to evaluate whether the free-water corrected metrics were more sensitive. As a last step, we evaluated associations between global Aβ SUVR and tau meta-ROI SUVR and white matter microstructure. Associations with a p-value < 0.05 were considered significant, but we also report associations that would survive false-discovery rate (FDR) correction for each bundle with q-value of 0.05, accounting for 6 tests (i.e. the number of diffusion measures assessed per bundle). Analyses were conducted using SPSS version 20 (IBM, N.Y., USA) and R version 3.6.3 (Vienna, Austria) (2020).

## 5 Acknowledgements

The authors wish to acknowledge the staff of PREVENT-AD as well as of the Brain Imaging Centre of the Douglas Mental Health University Institute and of the PET unit of the McConnell Brain Imaging Centre of the Montreal Neurological Institute, and members of the SCIL lab. A full listing of members of the PREVENT-AD Research Group can be found at https://preventad.loris.ca/acknowledgements/acknowledgements.php?date=[2020-06-30]. We would also like to acknowledge the participants of the PREVENT-AD cohort for dedicating their time and energy to helping us collect these data. Thank you to the Neuroinformatics Chair of the Université de Sherbrooke for supporting neuroscience research.

## Supplementary material

**TractQuerier query to extract the posterior cingulum**

~~~
import FreeSurfer.qry
#Posterior cingulum
Posterior_Cg.side = only(isthmuscingulate.side or posteriorcingulate.side and (entorhinal.side or fusiform.side or parahippocampal.side or precuneus.side or lingual.side or amygdala.side))
~~~

**Supplementary Figure 1.**
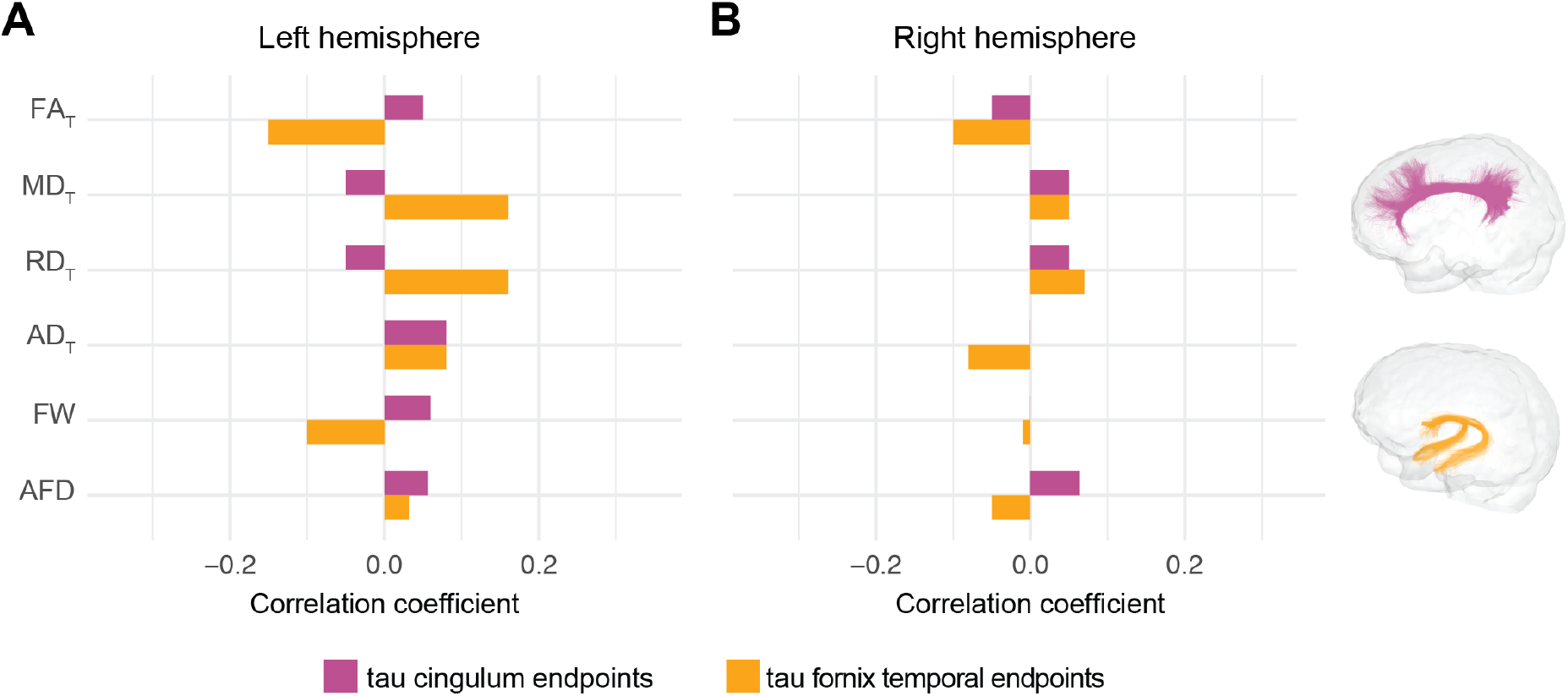
No associations between diffusion measures and tau in the cingulum and fornix. R_partial_ from regression models investigating associations between each diffusion metric (average diffusion measure in the bundle; independent variable) and tau pathology at the cortical endpoints of the bundle in the left (A) and right (B) hemispheres. Magenta bars correspond to associations in the cingulum and orange bars, in the fornix. Models included age, sex, bundle volume (divided by total intracranial volume) as covariates. Aβ: beta-amyloid; FAT: tissue fractional anisotropy; MD_T_: tissue mean diffusivity; ADT: tissue axial diffusivity; RD_T_: tissue radial diffusivity; FW: free-water index; AFD: fixel-based apparent fiber density

**Supplementary Figure 2.**
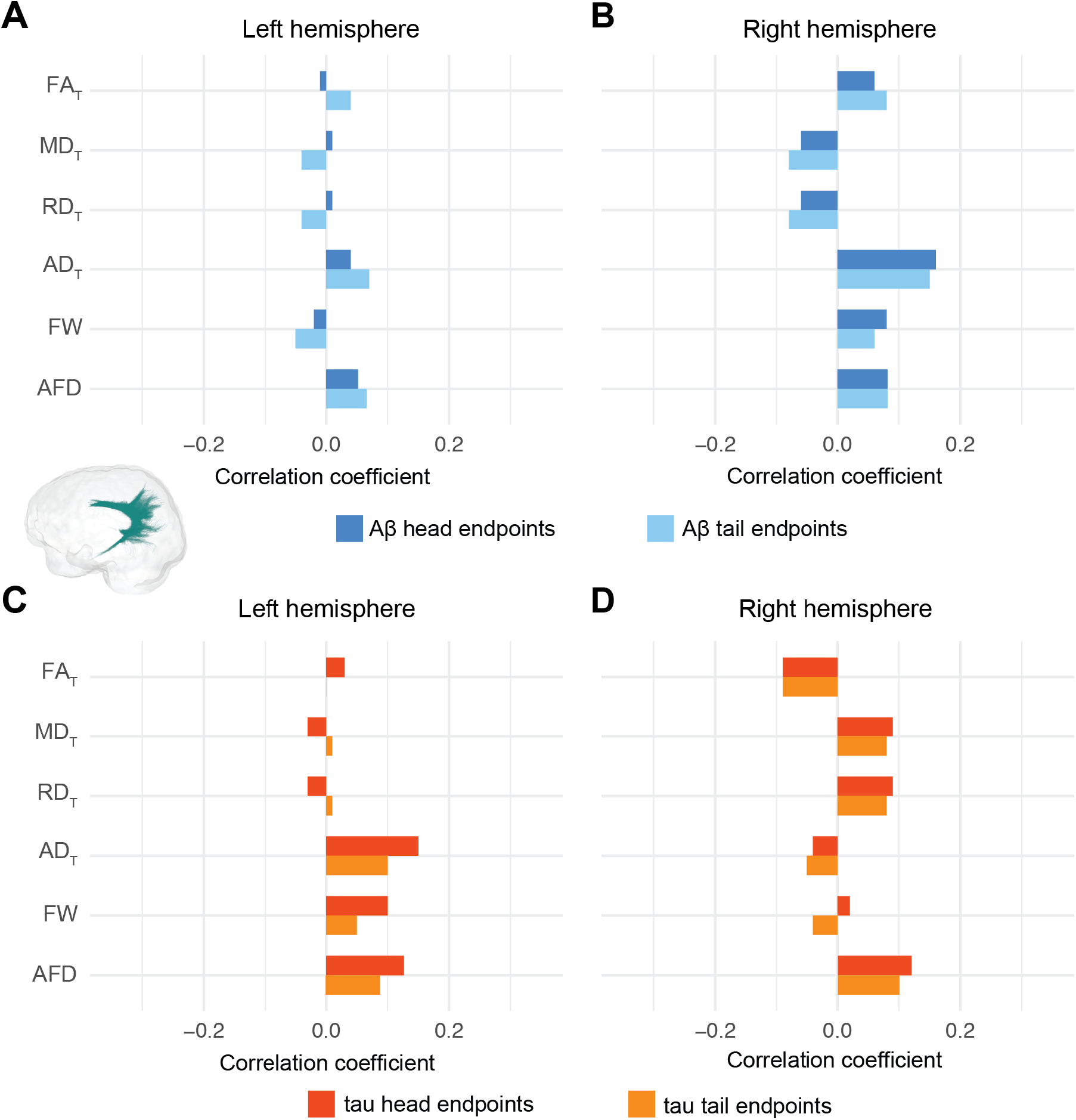
No associations between diffusion measures and pathology in the posterior cingulum. R_partial_ from regression models investigating associations between each diffusion metric (average diffusion measure in the bundle; independent variable) and Aβ (A-B) or tau (C-D) pathology at the cortical endpoints of the two ends of the posterior cingulum in the left hemisphere and right hemispheres. Models included age, sex, bundle volume (divided by total intracranial volume) as covariates. Aβ: beta-amyloid; FA_T_: tissue fractional anisotropy; MD_T_: tissue mean diffusivity; AD_T_: tissue axial diffusivity; RD_T_: tissue radial diffusivity; FW: free-water index; AFD: fixel-based apparent fiber density

**Supplementary Table 1.** Associations between diffusion measures and pathology in the uncinate fasciculus when adjusting for the other pathology.

| n=126 | Left hemisphere |  |  |  |  |  | Right hemisphere |  |  |  |  |  |
| --- | --- | --- | --- | --- | --- | --- | --- | --- | --- | --- | --- | --- |
| Associations with Aβ |  |  |  |  |  |  |  |  |  |  |  |  |
|  | FA<sub>T</sub> | MD<sub>T</sub> | AD<sub>T</sub> | RD<sub>T</sub> | FW | AFD | FA<sub>T</sub> | MD<sub>T</sub> | AD<sub>T</sub> | RD<sub>T</sub> | FW | AFD |
| Frontal endpoints | 0.16<br>(0.08) | -0.17<br>(0.07) | 0.07<br>(0.43) | -0.16<br>(0.07) | -0.08<br>(0.36) | 0.14<br>(0.14) | 0.13<br>(0.17) | -0.13<br>(0.17) | 0.13<br>(0.17) | -0.13<br>(0.17) | 0.06<br>(0.49) | 0.05<br>(0.57) |
| Temporal endpoints | 0.16<br>(0.08) | -0.16<br>(0.08) | 0.08<br>(0.38) | -0.16<br>(0.08) | 0.01<br>(0.96) | 0.11<br>(0.24) | 0.11<br>(0.24) | -0.11<br>(0.24) | 0.09<br>(0.31) | -0.11<br>(0.24) | 0.14<br>(0.12) | 0.00<br>(0.95) |
| Associations with tau |  |  |  |  |  |  |  |  |  |  |  |  |
| Frontal endpoints | 0.26<br>(0.00<br>4) | -0.26<br>(0.00<br>4) | 0.20<br>(0.03) | -0.26<br>(0.00<br>4) | -0.03<br>(0.74) | 0.19<br>(0.04) | 0.23<br>(0.01) | -0.23<br>(0.01) | 0.19<br>(0.03) | -0.23<br>(0.01) | 0.23<br>(0.01) | 0.14<br>(0.14) |
| Temporal endpoints | 0.16<br>(0.08) | -0.16<br>(0.08) | 0.14<br>(0.14) | -0.16<br>(0.08) | 0.05<br>(0.57) | 0.08<br>(0.41) | 0.19<br>(0.04) | -0.19<br>(0.04) | 0.17<br>(0.07) | -0.19<br>(0.04) | 0.27<br>(0.00<br>3) | 0.06<br>(0.55) |

R<sub>partial</sub> and (p-values) from regression models investigating associations between each diffusion metric (average diffusion measure in the bundle; independent variable) and pathology at the cortical endpoints of the two ends of the uncinate fasciculus (dependent variable). Models included age, sex, uncinate fasciculus volume (divided by total intracranial volume), and the amount of the other pathology at the corresponding endpoints as covariates. For example, in a model with Aβ at the frontal endpoints as the dependent variable, tau at the same endpoints is a covariate.

**Supplementary Table 2.** Associations between typical tensor measures and pathology in the uncinate fasciculus.

| n=126 | Left hemisphere |  |  |  | Right hemisphere |  |  |  |
| --- | --- | --- | --- | --- | --- | --- | --- | --- |
| Associations with Aβ |  |  |  |  |  |  |  |  |
|  | FA | MD | AD | RD | FA | MD | AD | RD |
| Frontal endpoints | 0.22<br>(0.02) | -0.13<br>(0.16) | 0.15<br>(0.11) | -0.20<br>(0.03) | 0.07<br>(0.46) | -0.01<br>(0.96) | 0.06<br>(0.53) | -0.04<br>(0.69) |
| Temporal endpoints | 0.14<br>(0.12) | -0.05<br>(0.63) | 0.14<br>(0.12) | -0.11<br>(0.22) | 0.01<br>(0.88) | 0.06<br>(0.55) | 0.08<br>(0.38) | 0.03<br>(0.73) |
| Associations with tau |  |  |  |  |  |  |  |  |
| Frontal endpoints | 0.28<br>(0.002) | -0.14<br>(0.13) | 0.20<br>(0.03) | -0.24<br>(0.01) | 0.12<br>(0.18) | 0.09<br>(0.32) | 0.24<br>(0.009) | 0.00<br>(0.98) |
| Temporal endpoints | 0.15<br>(0.11) | -0.02<br>(0.85) | 0.16<br>(0.08) | -0.09<br>(0.33) | 0.05<br>(0.60) | 0.17<br>(0.07) | 0.27<br>(0.003) | 0.08<br>(0.38) |

R<sub>partial</sub> and (p-values) from regression models investigating associations between each tensor metric (average diffusion measure in the bundle; independent variable) and pathology at the cortical endpoints of the two ends of the uncinate fasciculus (dependent variable). Models included age, sex, bundle volume (divided by total intracranial volume) as covariates.
Aβ : beta-amyloid; FA: fractional anisotropy; MD: mean diffusivity; AD: axial diffusivity; RD: radial diffusivity

